# Classifying Non-Small Cell Lung Cancer Histopathology Types and Transcriptomic Subtypes using Convolutional Neural Networks

**DOI:** 10.1101/530360

**Authors:** Kun-Hsing Yu, Feiran Wang, Gerald J. Berry, Christopher Ré, Russ B. Altman, Michael Snyder, Isaac S. Kohane

## Abstract

Non-small cell lung cancer is a leading cause of cancer death worldwide, and histopathological evaluation plays the primary role in its diagnosis. However, the morphological patterns associated with the molecular subtypes have not been systematically studied. To bridge this gap, we developed a quantitative histopathology analytic framework to identify the gene expression subtypes of non-small cell lung cancer objectively. We processed whole-slide histopathology images of lung adenocarcinoma (n=427) and lung squamous cell carcinoma patients (n=457) in The Cancer Genome Atlas. To establish neural networks for quantitative image analyses, we first build convolutional neural network models to identify tumor regions from adjacent dense benign tissues (areas under the receiver operating characteristic curves (AUC) > 0.935) and recapitulated expert pathologists’ diagnosis (AUC > 0.88), with the results validated in an independent cohort (n=125; AUC > 0.85). We further demonstrated that quantitative histopathology morphology features identified the major transcriptomic subtypes of both adenocarcinoma and squamous cell carcinoma (P < 0.01). Our study is the first to classify the transcriptomic subtypes of non-small cell lung cancer using fully-automated machine learning methods. Our approach does not rely on prior pathology knowledge and can discover novel clinically-relevant histopathology patterns objectively. The developed procedure is generalizable to other tumor types or diseases.

## Introduction

Non-small cell lung cancer accounts for 85% of lung cancer[1], with more than 1.4 million newly diagnosed patients per year worldwide[2,3]. Histopathology analysis by trained pathologists is the gold standard for diagnosing non-small cell lung cancer and defines the cancer types[4]. It is crucial to delineate lung malignancy from its morphologic mimic as specific treatment modalities (including surgical resection, chemotherapy, radiotherapy, and targeted therapy) can limit the progression of the disease and improve the survival outcomes of the patients[1]. In addition, the distinction between lung adenocarcinoma and squamous cell carcinoma, the two most common types of non-small cell lung cancer, is critical for selecting the optimal treatment: a few clinically actionable genetic variations[5,6] are almost exclusively observed in adenocarcinoma patients[4], whereas patients with squamous cell carcinoma respond better to gemcitabine[7] but could suffer from life-threatening hemoptysis when treated with bevacizumab[8,9]. Therefore, accurate histopathology diagnosis is crucial for formulating optimal treatment plans for lung cancer patients[10].

However, the current clinical process of histopathology assessment is not perfect[11,12]. Previous studies showed that there is a slight to moderate inter-observer variation in classifying malignant and benign lung tissues (κ=0.65-0.81)[13]. To estimate the diagnostic agreement for adenocarcinoma and squamous cell carcinoma, a group of researchers conducted an independent pathology review of 668 lung cancer cases and showed that the inter-observer agreement is moderate (κ=0.48-0.64)[11]. Another study suggested that the overall agreement for classifying adenocarcinoma and squamous cell carcinoma on morphologic criteria alone (in the absence of immunohistochemistry) was associated with expertise in pulmonary pathology (κ=0.41-0.46 among community pathologists, κ=0.64-0.69 among expert lung pathologists from the Pulmonary Pathology Society), and the diagnosis agreement did not reach the target for minimal clinical test reproducibility (κ=0.7) set by the investigators[12]. Further investigations on the histological patterns associated with lung adenocarcinoma subtypes showed that the overall inter-observer agreement (in kappa value) for stage IA tumors is 0.52, while that for stage IB tumors is 0.48[14]. Erroneous classification can lead to suboptimal treatment and loss of quality of life in patients.

In addition, the relations between histopathology morphology and the transcriptomics subtypes of lung adenocarcinoma and squamous cell carcinoma are not systematically studied. Transcriptomics subtypes of lung cancer are defined by the expression levels of a subset of genes related to the growth and differentiation of tumor cells[15,16]. It is not known whether the dysregulation of these key genes would impact the microscopic morphology of the tumor tissues. Through the systematic identification of transcriptomic-histopathology association, we can pinpoint the morphological changes linked with gene expression and dysregulation, thereby understand tumor cell morphology at the molecular level[5].

Computer vision algorithms, including convolutional neural networks, have shown exceptionally good performance for image classification[17]. These algorithms have demonstrated expert performance in several clinical domains including screening for diabetic retinopathy[18], identifying malignant dermatological lesions[19], and detecting cancer cells in pathology images[20,21]. With the recent availability of digital whole slide histopathology image in large cohorts[22,23], we can profile millions of tumor cells from a patient simultaneously and quantify the morphological differences in tumor cells among patients[24-26]. Leveraging terabytes of microscopic tissue image data, we can fine-tune the parameters of the neural networks to achieve optimal performance[27]. As the diagnosis of lung cancer is established by the morphological features of tumor cells and the diagnostic accuracy is positively associated with the evaluator’s experience[12], we hypothesize that convolutional neural networks trained on millions of histopathology image patches can distinguish malignancy from benign tissues, differentiate tumor types and identify the distinctive histopathology patterns of cancer cells. Moreover, the extensively documented transcriptomic distinctions between tumor subtypes[15,16,28] allow us to analyze the correspondence between transcriptomic differences and the morphologically-driven classifications. Here we used state-of-the-art computer vision methods to uncover the associations between transcriptomic subtypes and tumor cell morphology. The integrative transcriptomic-histopathology analysis will shred insight into the morphological changes related to molecular dysregulations.

In this study, we built convolutional neural network models to distinguish the histopathology and molecular subtypes of lung adenocarcinoma and squamous cell carcinoma. To ensure the generalizability of our methods, we validated the classification models in an independent cohort. We further identified the previously-unrecognized associations between tumor tissue morphology and transcriptomic profiles. Through this fully automated computational method, we can identify morphological differences in an unbiased fashion, which could be applied to provide decision support to clinicians encountering atypical histopathology changes[29,30], leverage quantitative morphology to study the macroscopic implications of transcriptomic patterns, and thereby contribute to precision cancer medicine[31,32].

## Methods

### Histopathology Images of Non-Small Cell Lung Cancer

Whole-slide histopathology images of lung adenocarcinoma (n=427) and lung squamous cell carcinoma (n=457) patients in The Cancer Genome Atlas (TCGA) cohort were obtained from the National Cancer Institute Genomic Data Commons[33,34]. To ensure the generalizability of our methods, an independent dataset from the International Cancer Genome Consortium (ICGC) cohort (87 lung adenocarcinoma and 38 lung squamous cell carcinoma patients) was acquired from the ICGC data portal[35]. All histopathology images were collected from primary, untreated tumors. Patients’ clinical profiles, such as age, gender, race, ethnicity, tumor stage, the anatomical subdivision of the neoplasm, as well as the accompanying pathology report were also obtained. The pulmonary pathologists’ evaluation from the TCGA and ICGC study consortiums were used as the ground truth for the diagnostic classification. Although a few samples in the TCGA dataset were collected more than ten years ago, the major diagnostic criteria for identifying tumor from benign tissues did not experience significant changes in the past decades[36]. The whole-slide images were broken into tiles with 1000×1000 pixels. Since the denser tiles contain more cells for further analysis, the 200 densest tiles for each whole slide were selected with the OpenSlide application programming interface (API)[37]. The resulting image tiles were rescaled for analysis by convolutional neural networks.

### Convolutional Neural Networks for Diagnosis Classification

Convolutional neural networks were built using the Caffe platform[38]. We evaluated several convolutional neural network implementations, including AlexNet[39], GoogLeNet[40], VGGNet-16[41], and the Residual Network-50 (ResNet)[42] because of their superior performance in prior image classification challenges[43]. AlexNet has a very efficient network design and employed non-saturating neurons to reduce training time[39]. The design of the GoogLeNet architecture is largely based on the Hebbian principle and has increased the depth and width of the network with a budgeted computational cost[40]. VGGNet possesses a deep and homogeneous convolution structure and demonstrates that the depth of a neural network is a crucial factor of its performance, and VGGNet-16 is the winner of the 2014 ImageNet Large Scale Visual Recognition Competition (ILSVRC) classification and localization task. [41]. ResNet is significantly deeper than VGGNet but lowered its model complexity by residual learning, and ResNet-50 is a 50-layer implementation of ResNet[42]. Classification models for histopathology images were fine-tuned from pre-trained ImageNet classification models based on these frameworks.

Three diagnostic classification tasks were performed: (1) to classify lung adenocarcinoma from adjacent dense benign tissues, (2) to classify lung squamous cell carcinoma from adjacent dense benign tissues, and (3) to classify lung adenocarcinoma from squamous cell carcinoma. The tiled images were the inputs to the classifier and the probabilities that the images belong to each category were the outputs of the convolutional neural network models.

### Evaluation of the Non-Small Cell Lung Cancer Diagnostic Classifiers

To evaluate the performance of the classifiers, the TCGA set was randomly divided into a training set (80% of the patients in the TCGA set) and a held-out test set (the remaining 20% of the patients). There was no overlap between patients in the training set and those in the test set, and the 80-20 split was guided by the machine learning literature[44]. The convolutional neural network models were trained and all hyper-parameters were finalized through cross-validation on the training set. Through this process, the optimized baseline learning rate was identified to be 0.001 for AlexNet and GoogLeNet, 0.0005 for VGGNet, and 0.01 for ResNet. Momentum was set to 0.9 and L2 regularization was used in all four architectures. The finalized models were first applied to the untouched TCGA test set and the predicted classification for each image was compared to the pathologists’ label. Receiver operating characteristics (ROC) curves of test set predictions were plotted and the areas under ROC curves (AUCs) were calculated. The AUCs of different classification tasks were compared. To ensure that the reported model performance on the TCGA held-out test set was not due to a fortuitous training-test set partition, the training-test set partition and the model training processes were repeated three times. Note that each time we retrained the model and re-optimized all parameters from scratch, to ensure the robustness of the results. The models trained and finalized by the TCGA training set were first validated by the TCGA test set and further evaluated by an independent validation set of histopathology images from the ICGC. The misclassified images by the machines were first examined by a physician-scientist (K.-H. Y.) in a blinded setting and independently reviewed by a pulmonary pathologist with more than 27 years of experience in lung pathology diagnosis (G. J. B.).

### Visualization of the Convolutional Neural Network Models

In order to interpret the convolutional neural network models, gradient-weighted class activation maps (Grad-CAM) were used to visualize the regions of importance in the classification process[45]. The Grad-CAM method identifies the gradient of the output from the pre-dense layer with respect to the input image, thereby characterizing the importance of the input pixels to the classification results[45]. For model visualization, the images with > 99.9% prediction confidence were retrieved and visualized. The Grad-CAM of models that distinguish lung adenocarcinoma from adjacent dense benign tissue, lung squamous cell carcinoma from adjacent dense benign tissue, and lung adenocarcinoma from lung squamous cell carcinoma were examined.

### Non-Small Cell Lung Cancer Transcriptomics Subtypes and Classifications

Previous research has described the transcription-based subtypes for both lung adenocarcinoma and lung squamous cell carcinoma[15,16]. Level 3 publicly-available gene expression data of the patient cohorts were acquired from the National Cancer Institute’s Genomic Data Commons and the methods described in the TCGA Consortium articles[33,34] were used to determine the transcriptomics subtypes of the patients. The samples with available gene expression data were randomly divided into an 80% training set and a 20% held-out test set[44]. The associations between histopathology patterns and the transcriptomic subtypes were investigated by building and fine-tuning a multi-class VGGNet convolutional neural network model using the training set. In the test phase, data from the held-out test set were inputted to the VGGNet model, and the output of the last layer of the VGGNet was obtained and transformed using principal component analysis, with the first two principal components visualized. In addition, pair-wise subtype classification models were trained on the subtypes with a sufficient number of cases (> 270 cases in the two subtypes combined) using the 80% training set, and the performance of the classification models was evaluated by their AUCs in the 20% held-out test set.

Patients with the same transcriptomic subtype nonetheless have variations in their transcriptomic signature. To evaluate the correlations between the variations in the transcriptomic signature that defined tumor subtypes and the histopathology-predicted subtypes in the test set, Spearman’s correlation coefficients were calculated between the transcriptomic subtype score[15,16] and the subtype probability predicted by a VGGNet model trained by histopathology images. The Spearman’s correlation test was performed to evaluate the strength of the correlations. All statistical analyses were performed in R version 3.3.

## Results

### Patient Characteristics

We obtained the whole-slide histopathology images from 427 lung adenocarcinoma[33] and 457 lung squamous cell carcinoma[34] patients in the TCGA database. Additional 87 patients with lung adenocarcinoma and 38 with lung squamous cell carcinoma were identified in the ICGC cohort. Pathology reports and clinical information of each patient, such as patient age, gender, race, and the anatomical subdivision of the tumor, were also acquired. Supplemental Table 1 summarizes the patient characteristics of the TCGA cohort, and Supplemental Table 2 outlines those for the ICGC cohort.

### Convolutional Neural Networks Classified Lung Adenocarcinoma from Adjacent Dense Benign Tissue

The convolutional neural network successfully distinguished lung adenocarcinoma from the adjacent dense benign tissue, with the areas under receiver operating characteristic curves (AUC) approximately 0.941-0.965 in the TCGA test set (VGGNet: 0.965 ± 0.012; ResNet: 0.952 ± 0.015; GoogLeNet: 0.965 ± 0.012; AlexNet AUC: 0.941 ± 0.014; Figure 1A). VGGNet and GoogLeNet performed slightly better than ResNet and AlexNet. The results were validated in the ICGC cohort, with AUCs 0.890-0.935 (VGGNet: 0.912 ± 0.037; ResNet: 0.935 ± 0.024; GoogLeNet: 0.901 ± 0.044; AlexNet AUC: 0.890 ± 0.051; Figure 1B). The performance on the ICGC dataset was not significantly different from that of the TCGA test set (Wilcoxon signed rank test P>0.15). Comparing with a previously-reported quantitative method for pathology analysis using the TCGA datasets[46], our neural networks attained 9-12% performance improvement in the TCGA test set. (Supplemental Table 3). We further investigated the gradient-weighted class activation maps (Grad-CAMs) and demonstrated that the convolutional neural networks give higher weights to regions indicative of tumorous changes (Figures 1C and 1D).

**Figure 1.**
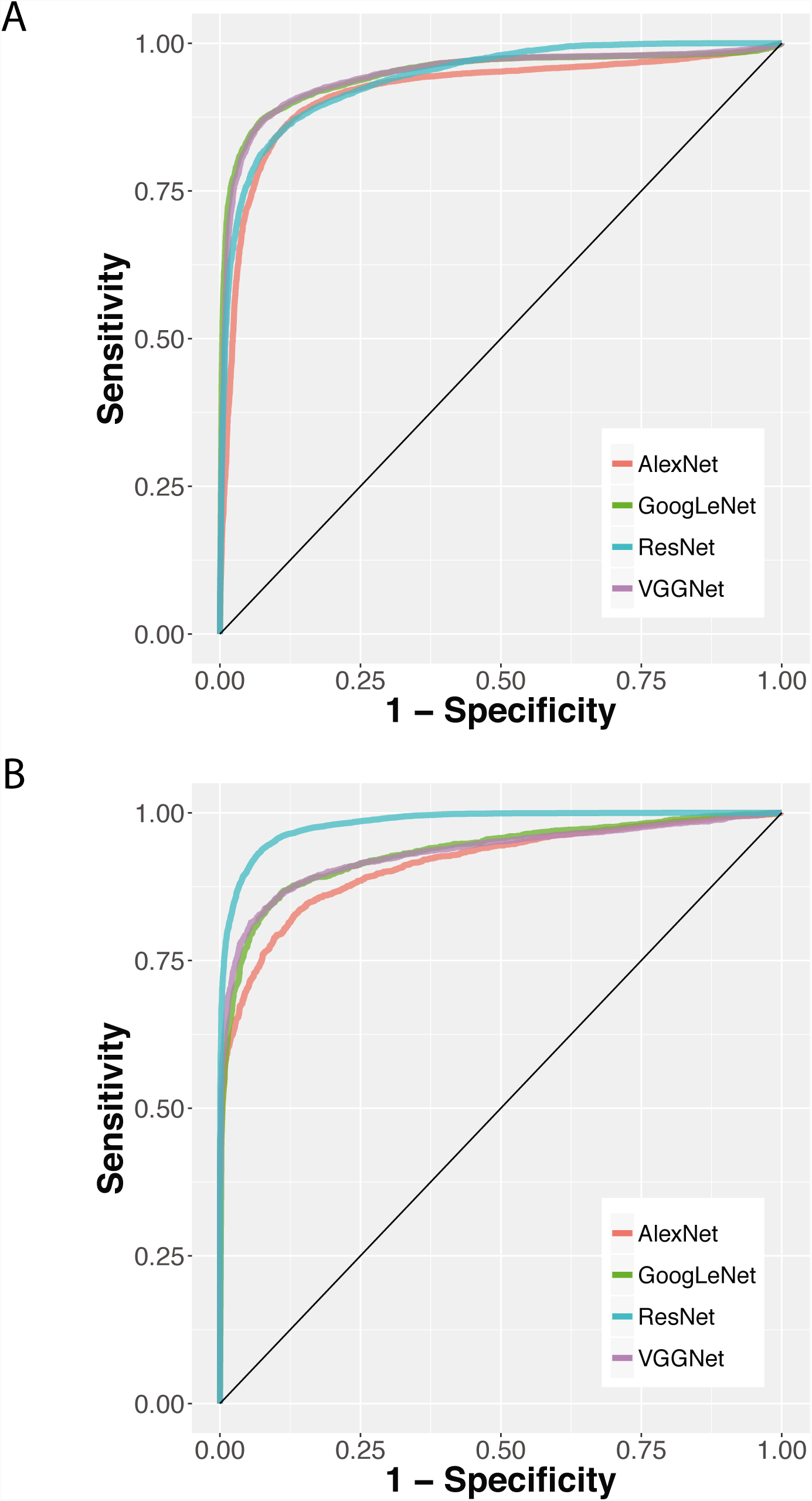
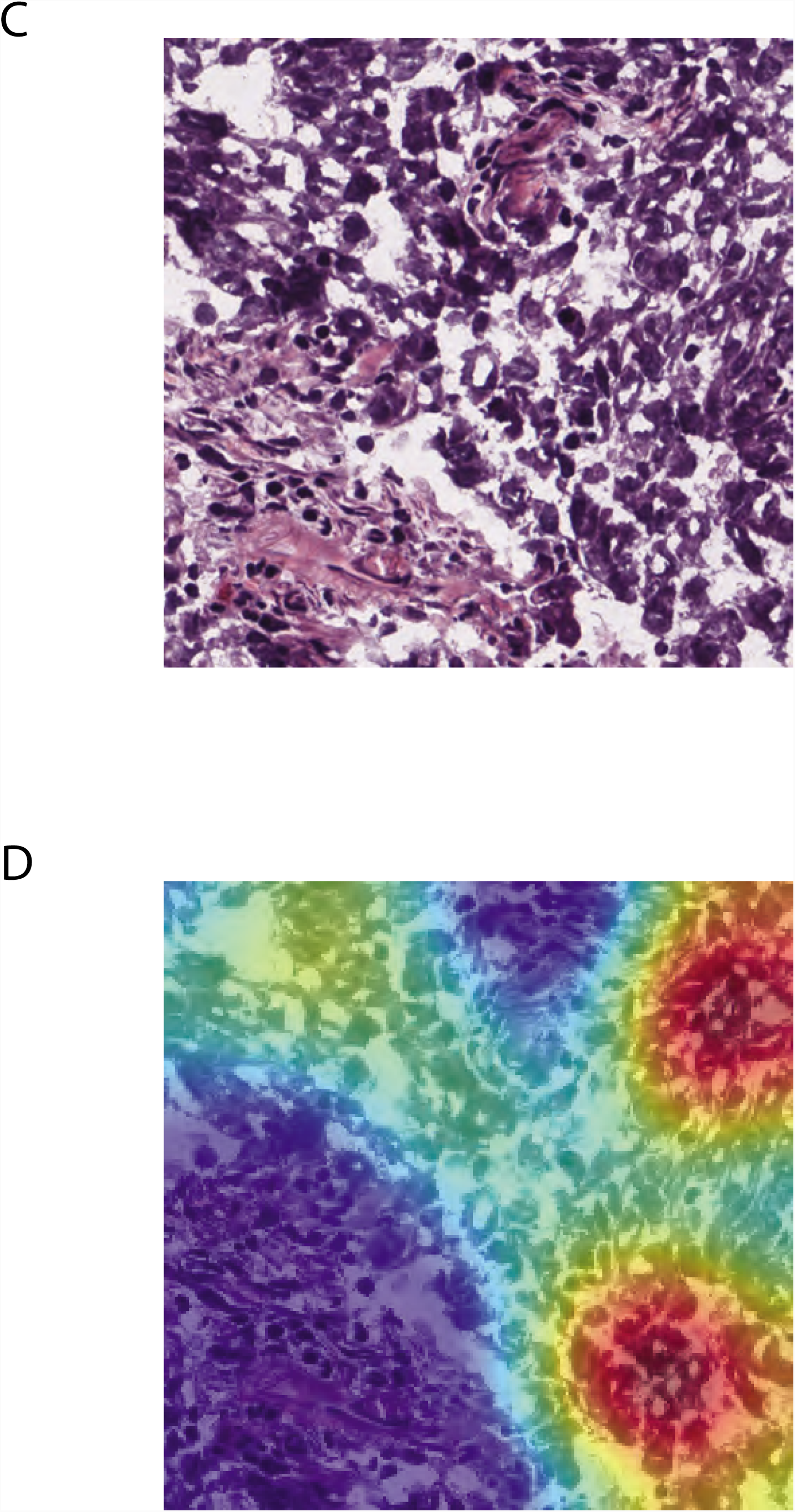
Convolutional neural networks distinguished lung adenocarcinoma from adjacent dense benign tissue, and the results were validated in the independent ICGC cohort. (A) Receiver operating characteristic (ROC) curves of AlexNet, GoogLeNet, ResNet, and VGGNet in the TCGA test set. Areas under receiver operating characteristic curves (AUCs): VGGNet: 0.965 ± 0.012; ResNet: 0.952 ± 0.015; GoogLeNet: 0.965 ± 0.012; AlexNet: 0.941 ± 0.014. (B) The ROC curves of the convolutional neural networks in the independent ICGC cohort. The classifiers achieved similar performance in this independent test set. AUCs: VGGNet: 0.912 ± 0.037; ResNet: 0.935 ± 0.024; GoogLeNet: 0.901 ± 0.044; AlexNet: 0.890 ± 0.051. (C) Attention analysis showed that the deep neural networks accurately utilized regions of tumorous changes to distinguish lung adenocarcinoma from adjacent dense benign tissues. A sample image tile of lung adenocarcinoma was shown. (D) Gradient-weighted class activation map (Grad-CAM) of the VGGNet model. The Grad-CAM method characterizes the regions where changes in pixel values would affect the classification score significantly, thereby quantifying where the artificial neural network put their “attention.” Regions with adenocarcinoma cells were identified and highlighted by the convolutional neural network automatically after training. Note that no human segmentation is involved in our training process.

### Convolutional Neural Networks Classified Lung Squamous Cell Carcinoma from Adjacent Dense Benign Tissue

Convolutional neural network classifiers achieved AUCs of 0.935-0.987 in distinguishing the tumor parts of lung squamous cell carcinoma from the adjacent dense benign tissue in the TCGA test set (VGGNet: 0.982 ± 0.0066; ResNet: 0.935 ± 0.0086; GoogLeNet: 0.987 ± 0.0047; AlexNet AUC: 0.977 ± 0.0040; Figure 2A). Similar performance was observed in the validation cohort from ICGC, with AUCs more than 0.979 (VGGNet: 0.991 ± 0.0019; ResNet: 0.979 ± 0.0035; GoogLeNet: 0.993 ± 0.0019; AlexNet AUC: 0.995 ± 0.0023; Figure 2B). Note that the performance on the ICGC dataset was not significantly different from that of the TCGA test set (Wilcoxon signed rank test P>0.15). In general, AlexNet, GoogLeNet, and VGGNet achieved similar classification performance, whereas the effectiveness of ResNet had slightly more variation. All of the neural network methods investigated achieved 6-11% increase in AUC, compared with previously reported machine learning methods developed with the TCGA datasets[46] (Supplemental Table 3). The high AUCs demonstrated the potential of identifying the suspicious part of the whole-slide images for further review and provide decision support to pathologists encountering ambiguous histopathology changes. Visualization of the attention map revealed that the convolutional neural networks attended to regions of squamous cancerous cell clusters, which validated the relevance of the classifiers (Figures 2C and 2D). Error analysis also revealed that images constantly misclassified by our models contained mislabels by TCGA investigators. Supplemental Figure 1 shows one of such images, which was tiled from a slide labeled as adjacent benign tissue by pathology evaluation conducted by TCGA, but a detailed pathology review by a pulmonary pathologist indicated that the frozen section image contains clusters of atypical cells and atypical glandular proliferation. Since the tissue slides were used for quality control purposes in the TCGA studies, the associated sample would need further evaluation in order to avoid biasing the omics profiling results in the original study.

**Figure 2.**
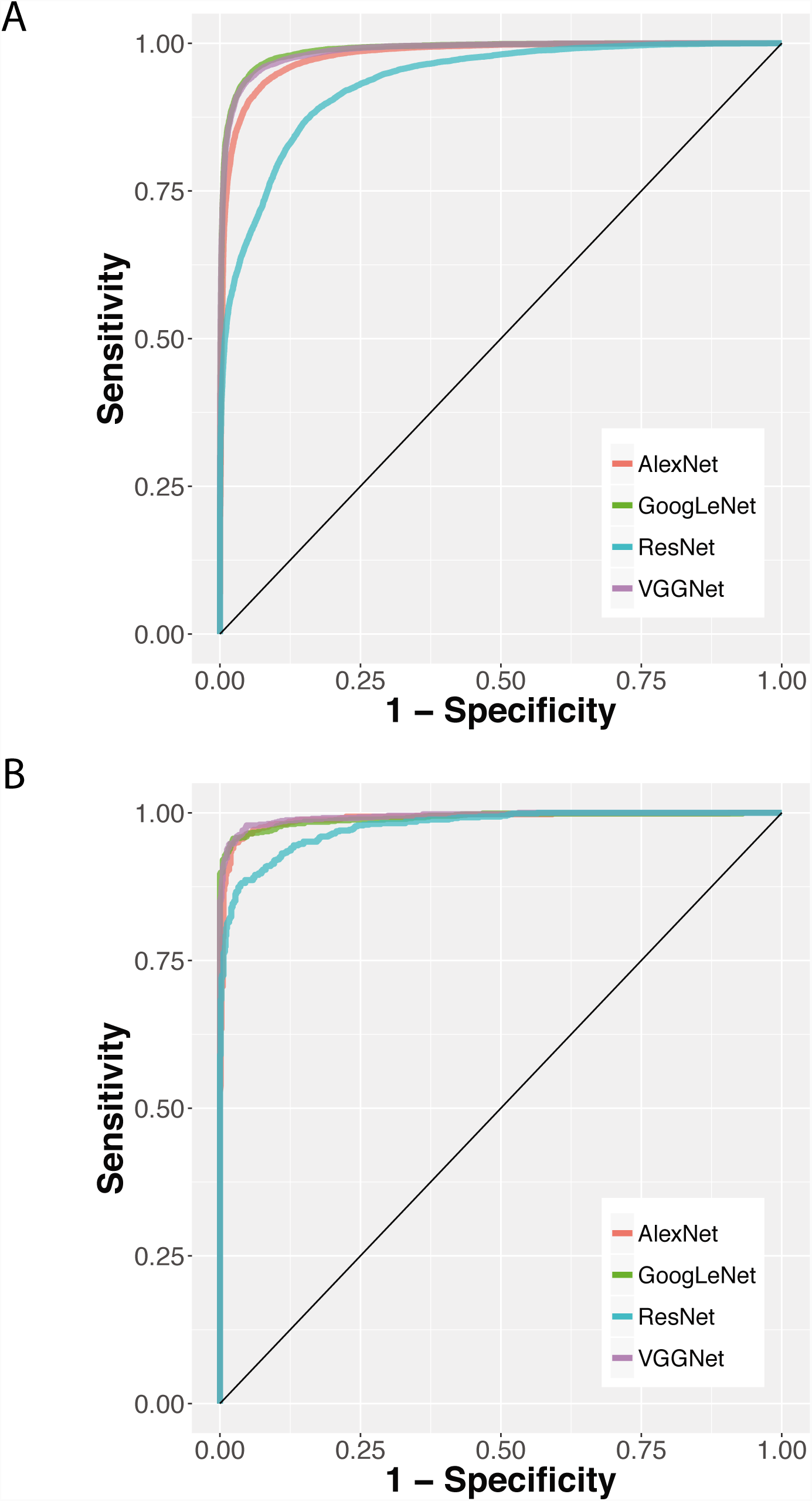
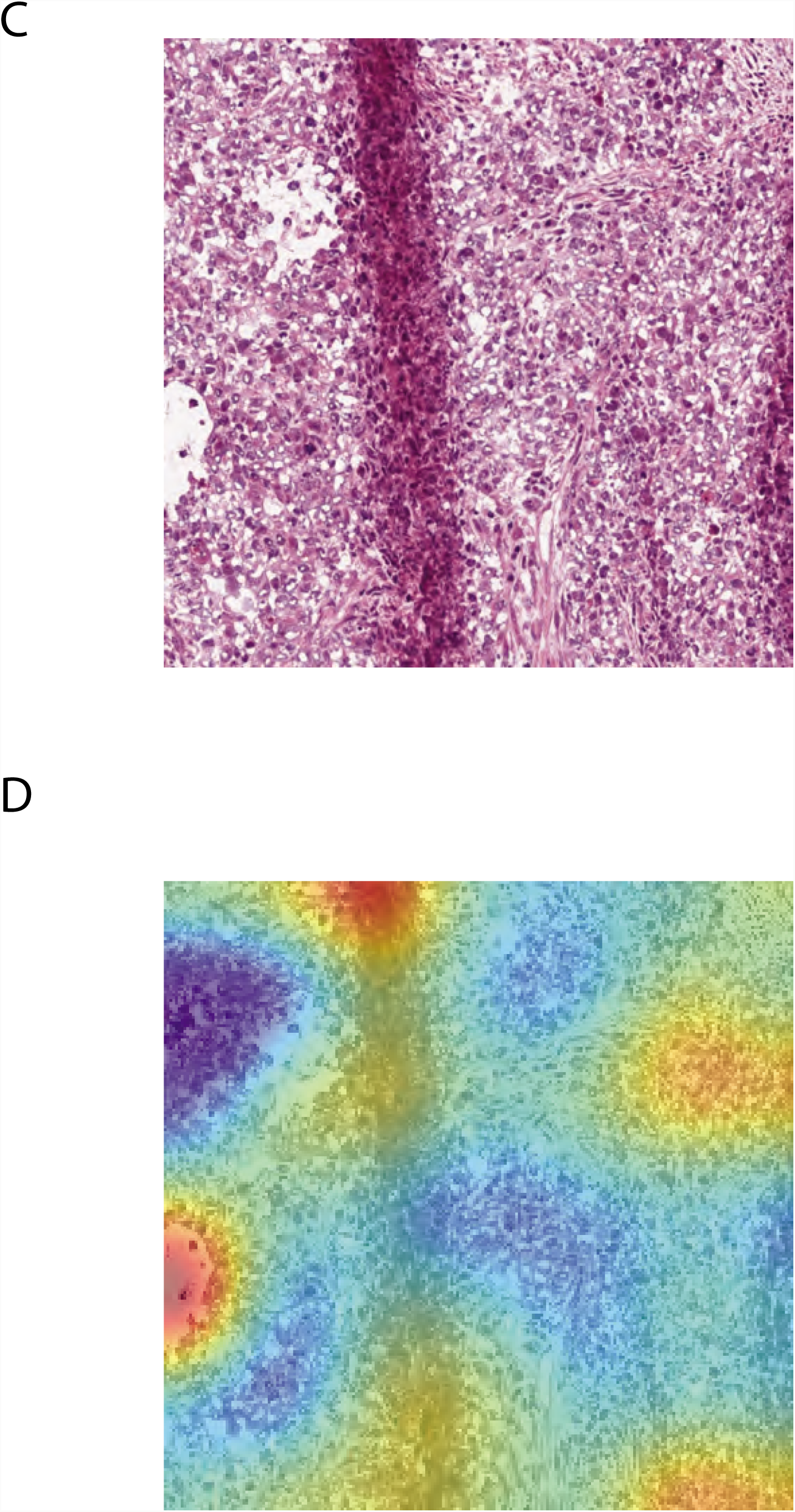
Convolutional neural networks distinguished lung squamous cell carcinoma from adjacent dense benign tissue, and the results were validated in the independent ICGC cohort. (A) Receiver operating characteristic (ROC) curves of AlexNet GoogLeNet, ResNet, and VGGNet in the TCGA test set. AUCs: VGGNet: 0.982 ± 0.0066; ResNet: 0.935 ± 0.0086; GoogLeNet: 0.987 ± 0.0047; AlexNet: 0.977 ± 0.0040. (B) The ROC curves of the convolutional neural network classifiers in the independent ICGC cohort. AUCs: VGGNet: 0.991 ± 0.0019; ResNet: 0.979 ± 0.0035; GoogLeNet: 0.993 ± 0.0019; AlexNet: 0.995 ± 0.0023. (C) Attention analysis of the deep neural networks demonstrated the regions of tumor cell clusters were used to distinguish lung squamous cell carcinoma from adjacent dense benign tissues. A sample image of lung squamous cell carcinoma was shown. (D) Gradient-weighted class activation map (Grad-CAM) of the VGGNet model. Regions of squamous cancer cells are automatically highlighted by the trained model, without human segmentation.

### Convolutional Neural Networks Classified Lung Adenocarcinoma from Lung Squamous Cell Carcinoma

Adenocarcinoma and squamous cell carcinoma are the two most common types of lung malignancy. It is crucial to distinguish them since the treatment options are different for these two cancer types, and the results demonstrated that convolutional neural networks achieved high accuracy in this task. The AUCs of the classifiers in the TCGA test set were approximately 0.883-0.932 (VGGNet: 0.932 ± 0.0042; ResNet: 0.883 ± 0.026; GoogLeNet: 0.930 ± 0.0013; AlexNet AUC: 0.896 ± 0.0226; Figure 3A). All of the neural network models performed 12-27% better than the feature-based machine learning methods on the TCGA test set[46] (Supplemental Table 3). The AUCs of the finalized models applied to the validation cohort (ICGC) were 0.752-0.857. (VGGNet: 0.843 ± 0.019; ResNet: 0.857 ± 0.024; GoogLeNet: 0.830 ± 0.014; AlexNet AUC: 0.752 ± 0.016; Figure 3B). The performance on the ICGC dataset was not significantly different from that of the TCGA test set (Wilcoxon signed rank test P>0.09). Except for the varying performance of AlexNet, the neural network with the simplest architecture among the four, the performance of the other neural network frameworks was comparable in both cohorts. Grad-CAM analyses revealed the distinctive visual patterns of the two tumor types picked up by the convolutional neural networks, such as the clustering patterns of tumor cells (Figures 3C-3F).

**Figure 3.**
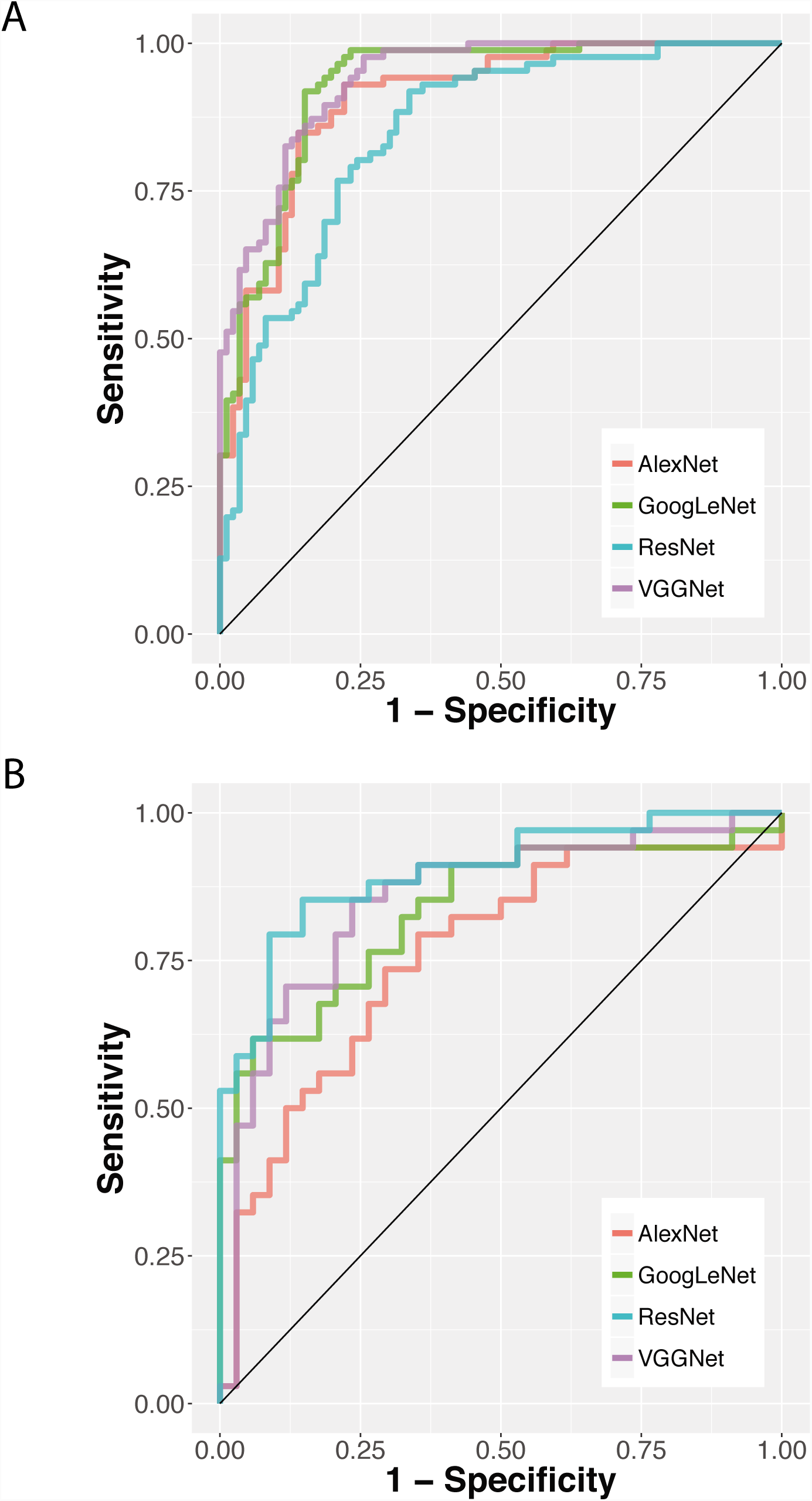
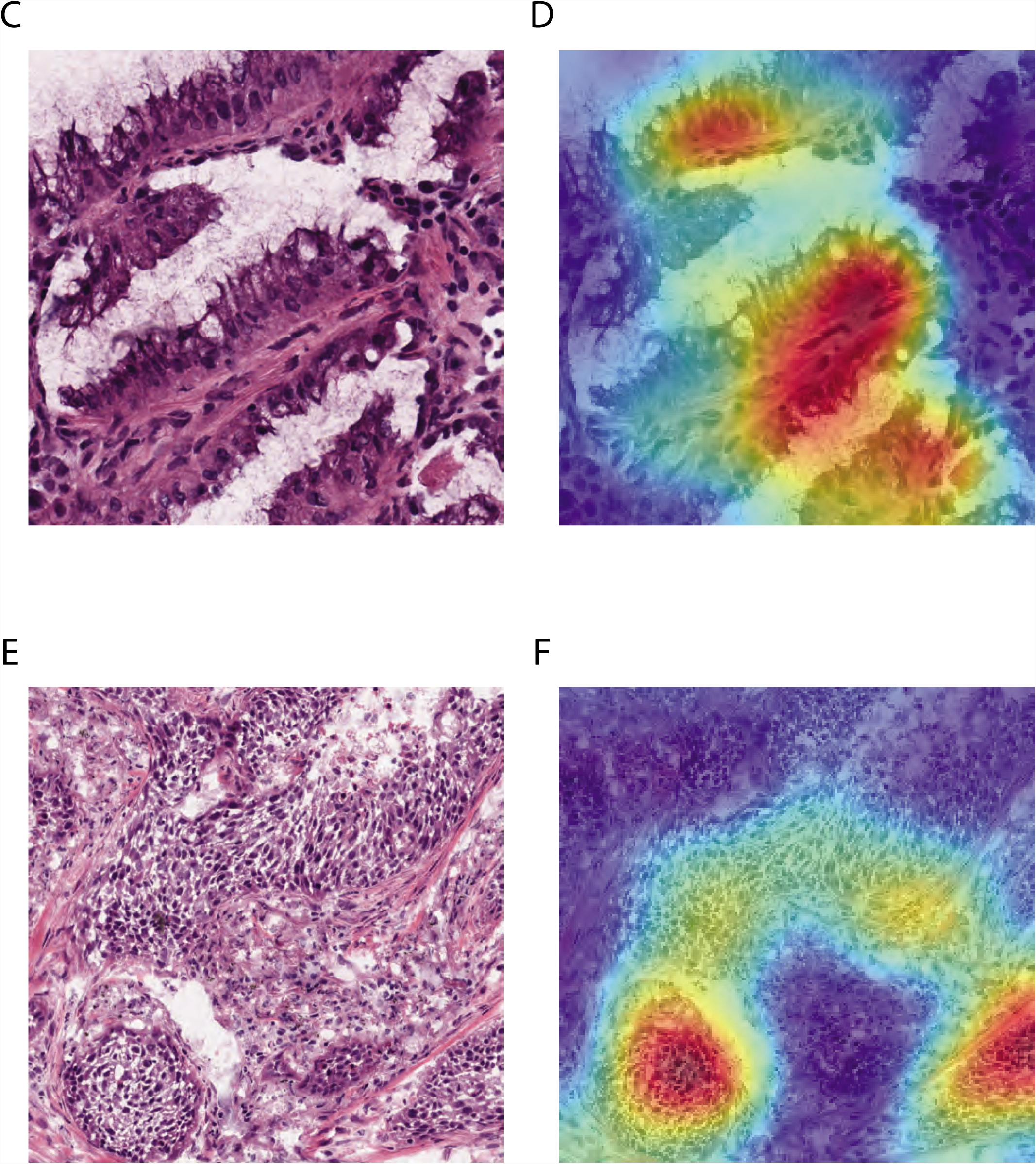
Convolutional neural networks distinguished lung squamous cell carcinoma from lung adenocarcinoma, and the results were validated in the independent ICGC cohort. (A) Receiver operating characteristic (ROC) curves of AlexNet, GoogLeNet, ResNet, and VGGNet in the TCGA test set. AUCs: VGGNet: 0.932 ± 0.0042; ResNet: 0.883 ± 0.026; GoogLeNet: 0.930 ± 0.0013; AlexNet: 0.896 ± 0.0226. (B) The ROC curves of the convolutional neural network classifiers in the independent ICGC cohort. AUCs: VGGNet: 0.843 ± 0.019; ResNet: 0.857 ± 0.024; GoogLeNet: 0.830 ± 0.014; AlexNet: 0.752 ± 0.016. (C) Visualization of the attention map showed regions the deep neural networks attended to when distinguishing lung squamous cell carcinoma from lung adenocarcinoma. A sample image of lung adenocarcinoma was shown. (D) The Grad-CAM plot of the lung adenocarcinoma image. (E) A sample image of lung squamous cell carcinoma. (F) The Grad-CAM plot of the lung squamous cell carcinoma image. Note that no human segmentation is required in our training process.

### Convolutional Neural Networks Correlated Cell Morphology with Gene Expression Subtypes

Previous studies defined the subtypes of non-small cell lung cancer by the clusters of tumor gene expression profiles. The three transcriptomic subtypes of lung adenocarcinoma proposed by TCGA are the terminal respiratory unit (TRU), proximal inflammatory (PI), and proximal proliferative (PP) subtypes, which correspond to the previous pathological, anatomic, and mutational classifications of bronchioid, squamoid, and magnoid subtypes respectively[16,33]. The four transcriptomic subtypes of lung squamous cell carcinoma proposed by TCGA are the classical, basal, secretory, and primitive subtypes[15,34]. These subtypes are related to the differentially activated inflammation, proliferation, and cell differentiation pathways[15,16]. Here we analyzed the lung adenocarcinoma and squamous cell carcinoma patients with available histopathology slide and RNA-sequencing data (Supplemental Table 4), determined their transcriptomic subtypes, and employed convolutional neural networks to associate the morphological patterns of the cells with patients’ molecular subtypes. A multiclass image classification model showed that the principal components of the output values from the last layer of VGGNet were significantly correlated with the transcriptomic subtypes of both adenocarcinoma (ANOVA P < 0.001 in both principal components 1 and 2; Figure 4A) and squamous cell carcinoma (ANOVA P < 0.001 in principal component 1; Figure 4C). Correlation analysis showed that patients with typical transcriptomic profiles for the subtype they belong to also possessed histopathology patterns typical of that subtype (Figures 4B and 4D). The Spearman’s correlation coefficients of the transcriptomics-based subtype scores and the histopathology-derived probabilities were greater than 0.4 in all three subtypes (P<0.01) of lung adenocarcinoma and in the two common subtypes (classical and prototypical) of lung squamous cell carcinoma. In addition, pair-wise neural network models predicted the subtypes of lung adenocarcinoma with AUCs 0.771-0.892 in the best classifiers (Supplemental Figure 2) and predicted the major subtypes of lung squamous cell carcinoma with AUCs approximately 0.7 (Supplemental Figure 3). The slightly worse performance in squamous cell carcinoma might originate from the fact that there were fewer numbers of samples in each of the four subtypes. These results indicated the strong correlations between cell morphology and gene expression subtypes of non-small cell lung cancer.

**Figure 4.**
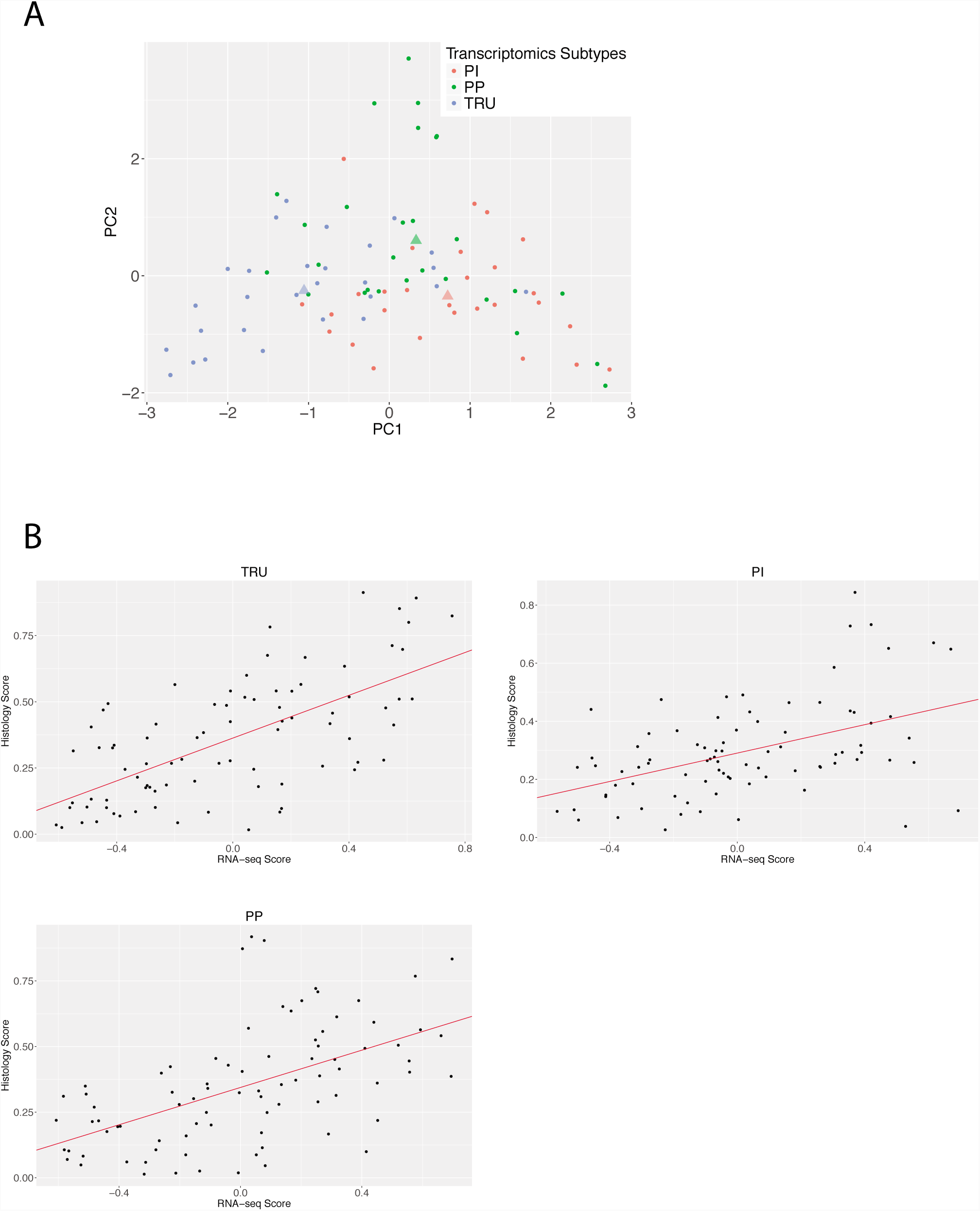
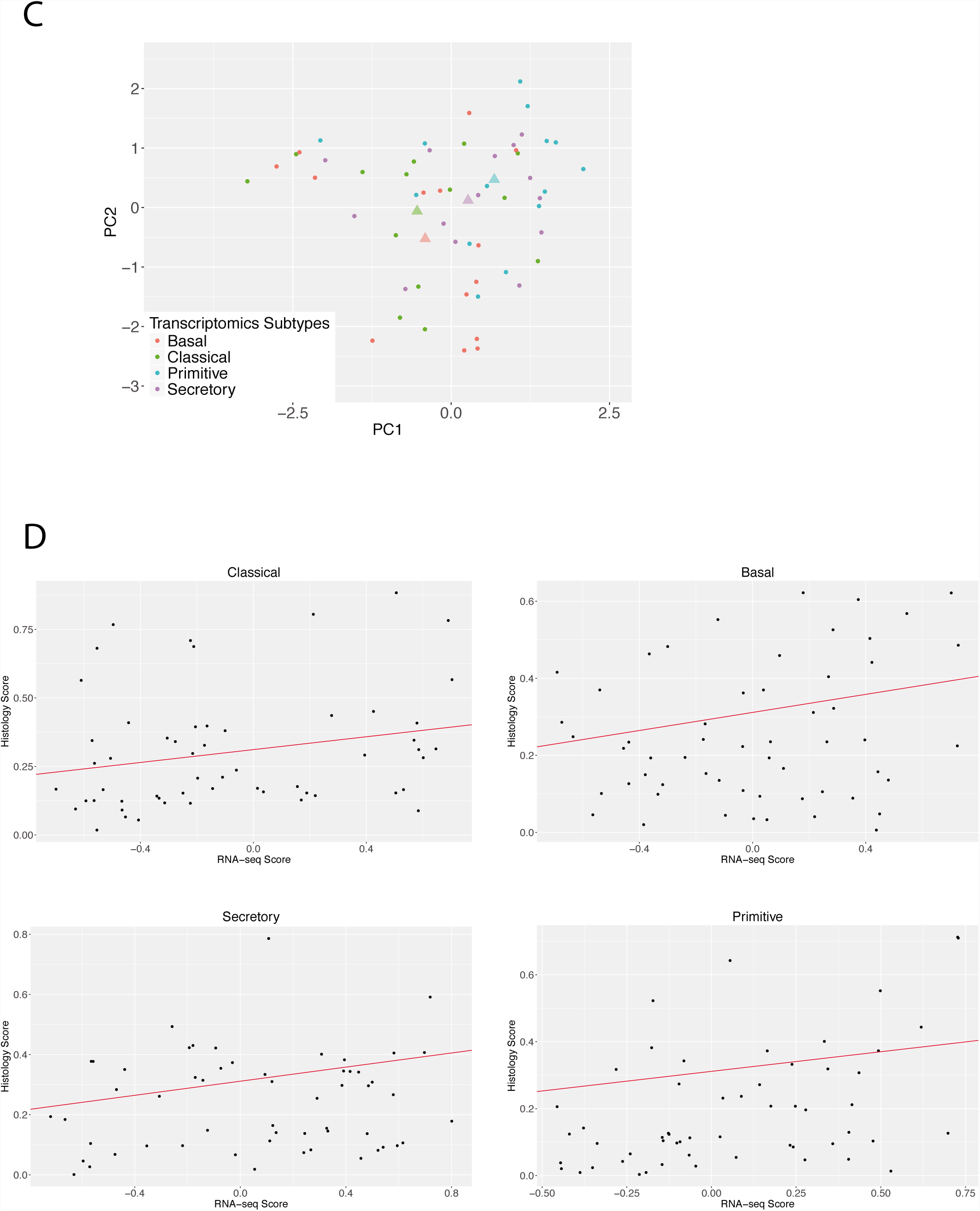
Convolutional neural network associated transcriptomic subtypes with histopathology patterns. (A) The principal components (PCs) of the VGGNet-based histopathology image summary vector were correlated with the three subtypes of lung adenocarcinoma (PC1 ANOVA P-value < 0.0001; PC2 ANOVA P-value = 0.0038). Each dot represents a sample in the test set. The color of the dots indicates the transcriptomic subtypes of the sample. The triangles represent the mean PC1 and PC2 of each subtype. TRU: terminal respiratory unit subtype; PI: proximal inflammatory subtype; PP: proximal proliferative subtype. (B) Correlation plots of the transcriptomic subtype scores and the histopathology classification scores in lung adenocarcinoma. Significant associations were found in all three subtypes (P<0.01). (C) The first principal component of the VGGNet-based histopathology image summary vector is correlated with the four subtypes of lung squamous cell carcinoma (PC1 ANOVA P-value = 0.0028). Each dot represents a sample in the test set. The color of the dots indicates the transcriptomic subtypes of the sample. The triangles represent the mean PC1 and PC2 of each subtype. (D) Correlation plots of the transcriptomic subtype scores and the histopathology classification scores in lung squamous cell carcinoma. Significant associations were found in the classical and primitive subtypes (P<0.01).

## Discussion

This is the first study that used convolutional neural networks to identify the transcriptomic subtypes of lung malignancy. Results demonstrated that the deep learning framework identified histopathology slides with tumor cells, captured the cell morphologies related to lung cancer diagnosis, and correlated histopathology with transcriptomic profiles. The diagnostic classifications were successfully validated in an independent test set, showing the robustness of our methods. Our analytical framework requires no human intervention after tissue slide preparation, contributes to building an automated decision support system, and points to the associations between tumor cell morphology and molecular biology.

Our work demonstrates a systematic approach to analyzing histopathology images in a non-biased fashion. Compared with the previous methods[24-26,46], our approaches require no human segmentation[24] and feature definition,[24-26,46] making them easily fit into the clinical workflow of pathology diagnosis. The extracted features do not rely on prior pathology knowledge. This enables us to identify novel morphological patterns related to clinically relevant phenotypes and biological processes at the gene expression level. In addition, our models performed 6-27% better than the previously-proposed feature-based methods, and the error rates were comparable with the reported inter-observer variations, indicating the utility of deep convolutional neural network in classifying lung cancer types. Furthermore, our system successfully identified regions with atypical cells and atypical glandular proliferation in slides that were marked as adjacent dense benign tissue by TCGA (Supplemental Figure 1). This finding was confirmed by an experienced pulmonary pathologist. This suggests that our methods could learn the general patterns of pathology in the presence of noise and mislabels in the dataset, which could identify suspicious cells in the histopathology slides of research samples and provide decision support for pathology evaluation[29,47]. Given the increasing incidence rate of lung cancer and the projected shortage of pathologists[48], we can augment the current pathology evaluation workflow with the reported system, which can double-check on the diagnoses made by human practitioners and point out suspicious tissue regions requiring additional review. The deployment of machine-learning systems in the clinical settings has the potential of reducing the cost and loss of quality of life associated with misdiagnoses[49]. In addition, the classification of transcriptomic subtypes can facilitate further studies on the morphological impact of aberrant gene expression in the tumor tissue.

The general architecture of our convolutional neural networks was trained on the images from the ImageNet[43], which bear little resemblance to histopathology images. However, we showed that these frameworks generated good classification performance when refined with an adequate amount of training data for model fine-tuning[50]. This indicates the utility of fine-tuning the pre-trained convolutional neural networks using whole-slide pathology training images from hundreds of patients. When classifying cancerous regions from the adjacent dense benign tissues, AlexNet, which consists of only 5 convolutional layers, has similar performance compared with GoogLeNet, VGGNet, and ResNet. However, for more sophisticated tasks, such as differentiating tumor types, AlexNet has the worst performance among all convolutional neural networks we investigated. These results indicated that simple models may be suitable for tumor detection tasks, while diagnosing tumor types and subtypes may require neural networks with more layers.

As expected, convolutional neural networks worked best when the number of training samples was large[51,52]. Whole slide histopathology images of tumor tissues provide a substantial opportunity for convolutional neural network applications, as one slide typically contains thousands of tumor cells. In addition, several types of tumor cell variations are often represented in different parts of the same whole-slide image. The abundance of tumor cells and their variations per slide provided sufficient data for establishing a diagnostic model for non-small cell lung cancer. This study demonstrated the feasibility of developing a deep learning-based platform for histopathology diagnosis and omics-histopathology integration. The developed methods are extensible to other imaging classification tasks central to clinical diagnosis[53]. Further studies are needed to establish the cell-level image-transcriptomics association by single-cell sequencing and validate the utility of computer vision methods for diagnosing other prevalent tumor types.

One limitation of the study is that the TCGA and ICGC datasets only included the two most common histology types of lung cancer: adenocarcinoma and squamous cell carcinoma. Although these two subtypes comprise more than 80% of non-small cell lung cancer, the diagnostic performance of rarer types of tumors, such as giant-cell carcinoma of the lung or small cell lung carcinoma, cannot be evaluated based on the available data[54,55]. Large-scale collection of rarer cancer types and prospective studies on the efficacy of the developed models are required to enable comprehensive cancer diagnostic systems[49]. In addition, all of the histopathology images in this study were gathered retrospectively, and the updated International Association for the Study of Lung Cancer/American Thoracic Society/European Respiratory Society classifications of lung adenocarcinoma[56] were not available for the study cohorts. Future studies are needed to quantify the inter-observer disagreement of various lung cancer pathology and investigate the clinical utility of implementing an automated histopathology image analysis system in real-world settings[57].

Overall our study demonstrated the utility of convolutional neural networks in associating tumor cell morphology with their molecular subtypes and classifying the histopathology images of the major types of non-small cell lung cancer. The machine learning system presented here can provide decision support to pathologists, detect atypical cells in large datasets with noisy labels, and aid in reclassifying patients with inconclusive histopathology presentations. Further testing in the clinical settings is needed to confirm the utility of our system. Our developed bioinformatics workflow is generalizable to other tumor types or diseases.

## Acknowledgments

The authors would like to thank Dr. Matt van de Rijn for his valuable advice on the study design and interpretation of results, and Ce Zhang, Alex Ratner, Andrej Krevl, and Rok Sosic for their assistance on the computation framework. We thank the AWS Cloud Credits for Research, Microsoft Azure for Research Award, and the NVIDIA GPU Grant Program, and the Extreme Science and Engineering Discovery Environment (XSEDE) at the Pittsburgh Supercomputing Center (allocation TG-BCS180016) for their computational supports. We thank the anonymous reviewers for their constructive feedback.

## Author Contributions

K.-H.Y. conceived, designed, performed the analyses, interpreted the results, and wrote and revised the manuscript. F.W. implemented an early version of neural networks and edited the manuscript. G.J.B., C.R., R.B.A., M.S, and I.K. interpreted the results and edited the manuscript. I.K. supervised the work. K.-H.Y. had full access to all the data in the study and takes responsibility for the integrity of the data and the accuracy of the data analysis.

## Figure Legends

**Supplemental Figure 1.** Error analysis revealed that images constantly misclassified by convolutional neural network models were mislabeled by TCGA. An example image tile labeled as adjacent benign tissue by TCGA contains clusters of atypical cells and atypical glandular proliferation.

**Supplemental Figure 2.** Convolutional neural networks identified the correlations between histopathology image patterns and three transcriptomic subtypes of lung adenocarcinoma. The AUC for pair-wise classification is greater than 0.77 in the best classifiers. (A) The ROC curves of classifying images of the terminal respiratory unit subtype from those of the proximal inflammatory subtype, with AUCs 0.816-0.892. (B) The ROC curves of classifying images of the terminal respiratory unit subtype from those of the proximal proliferative subtype, with AUCs 0.771-0.867. (C) The ROC curves of classifying images of the proximal inflammatory subtype from those of the proximal proliferative subtype, with AUCs 0.687-0.771.

**Supplemental Figure 3.** Histopathology image patterns correlated with the prevalent transcriptomic subtypes of lung squamous cell carcinoma. The AUC is approximately 0.70. (A) The ROC curves of classifying images of the classical subtype from those of the basal subtype. VGGNet is the best classifier, with AUC 0.685 ± 0.018. (B) The ROC curves of classifying images of the classical subtype from those of the secretory subtype. VGGNet is the best classifier, with AUC 0.701 ± 0.061. The reduced AUC might result from the smaller sample size of the four subtypes of lung squamous cell carcinoma in our cohorts.

