## Supplementary figures and images for "Classifying Non-Small Cell Lung Cancer Histopathology Types and Transcriptomic Subtypes using Convolutional Neural Networks"

### Supplemental Figure 1, Supplemental Figure 2, Supplemental Figure 3

Supplemental Figure 1

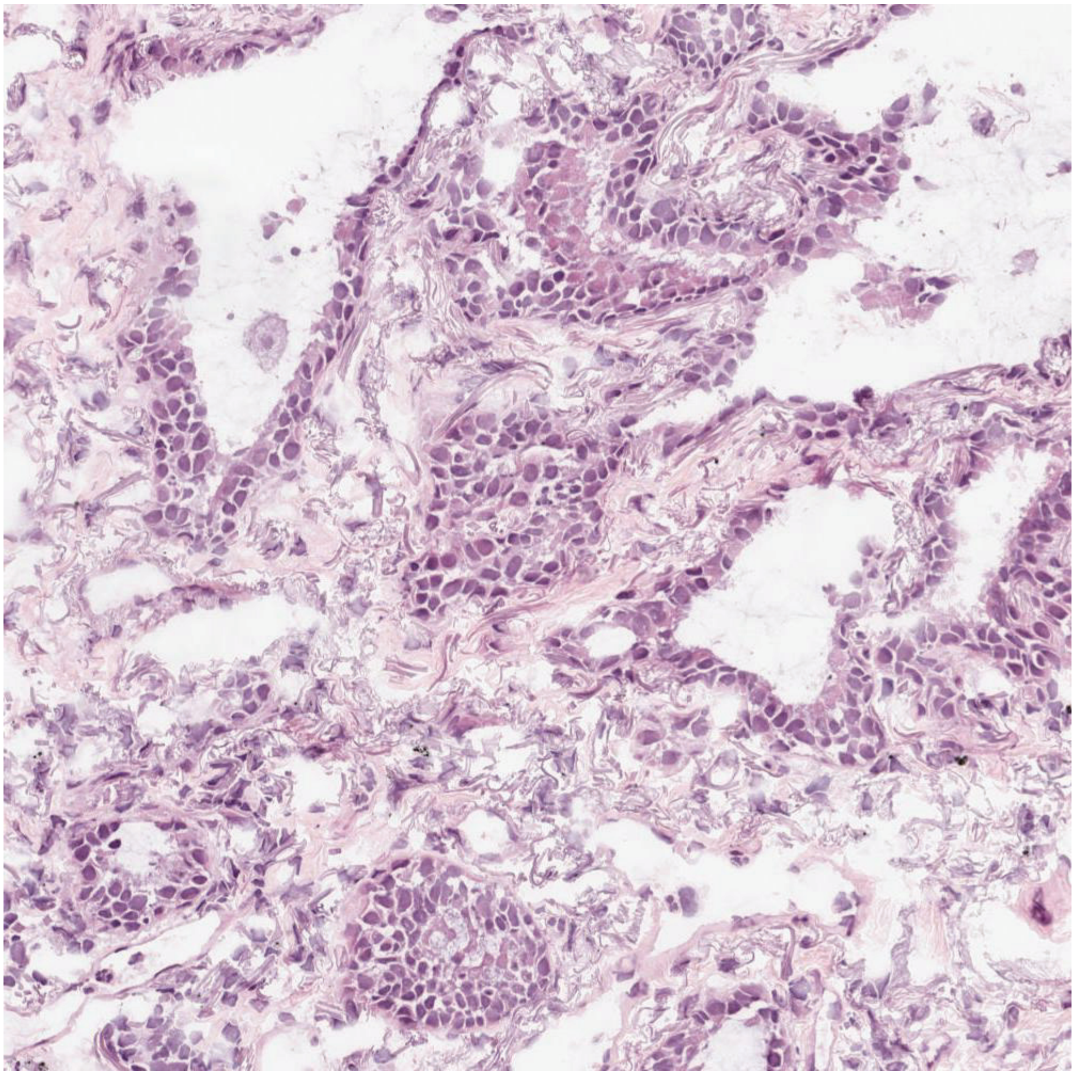

#### Supplemental Figure 2

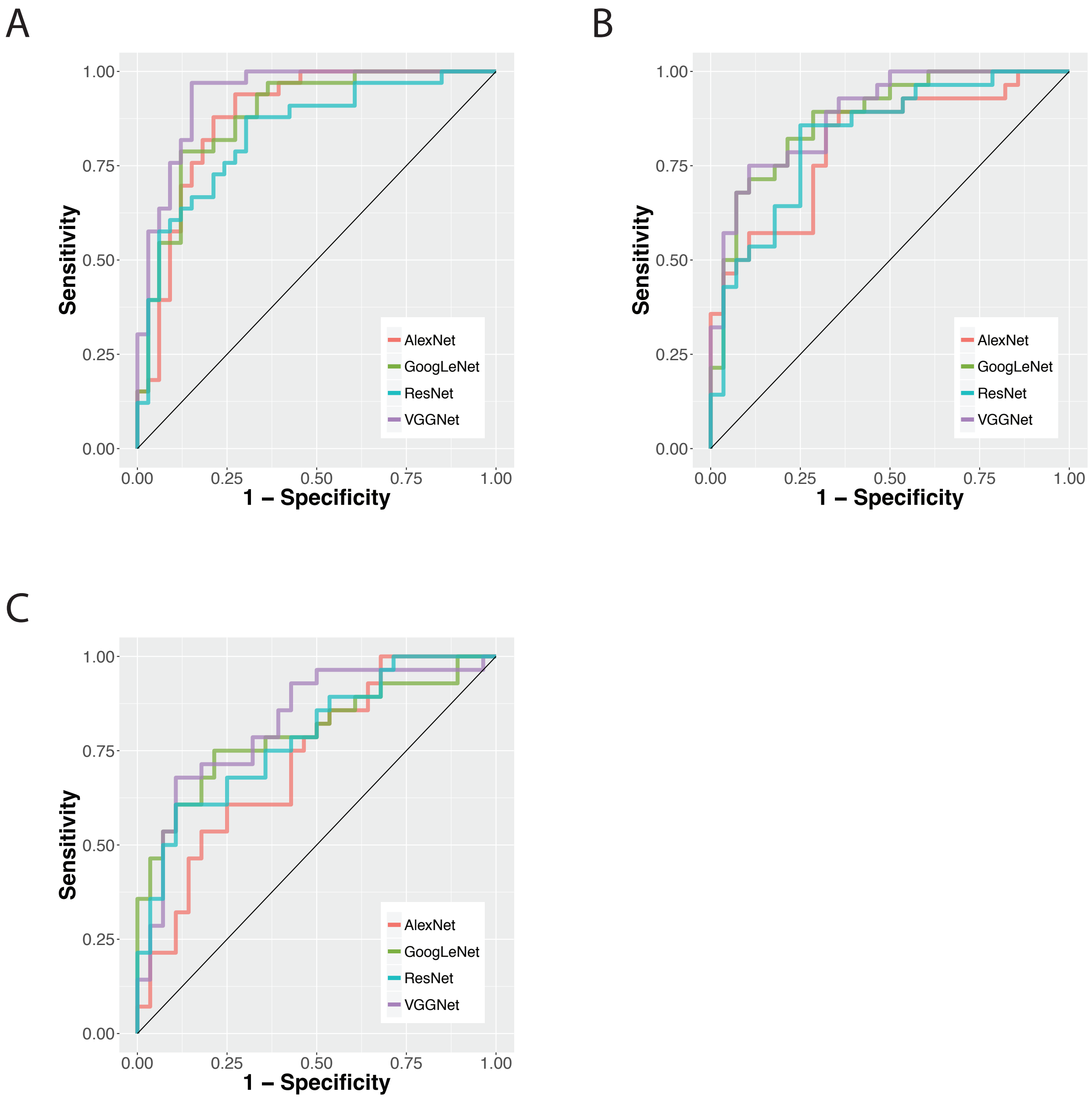

Supplemental Figure 3

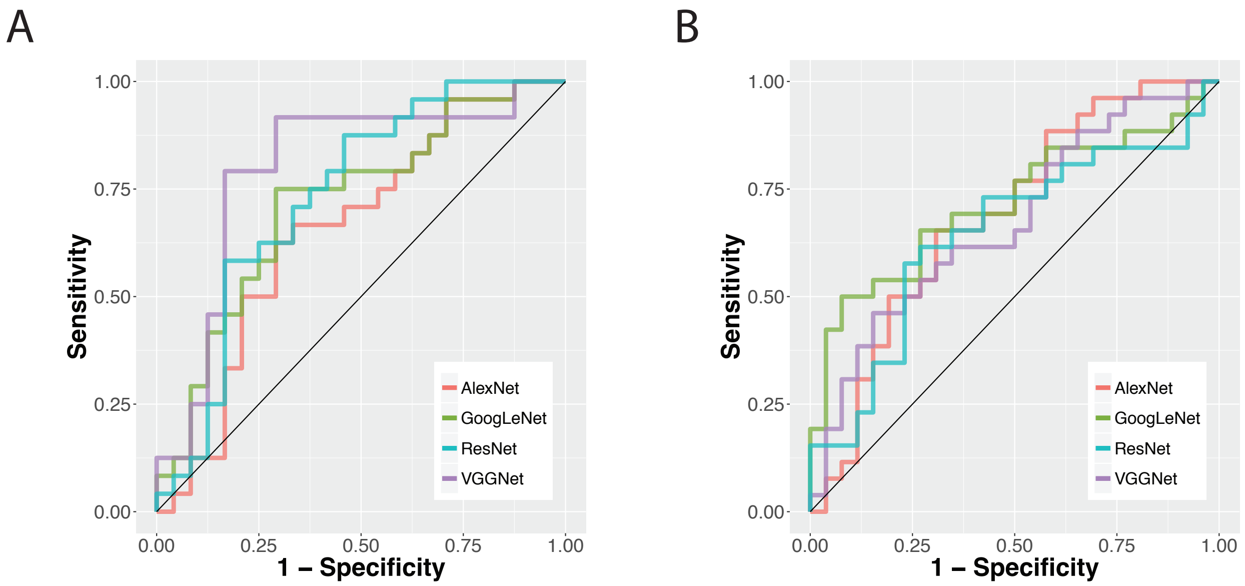
