## Supplemental Table 1, Supplemental Table 2, Supplemental Table 3, Supplemental Table 4 for "Classifying Non-Small Cell Lung Cancer Histopathology Types and Transcriptomic Subtypes using Convolutional Neural Networks"

**Supplemental Table 1.** Characteristics of lung adenocarcinoma and lung squamous cell carcinoma patients in The Cancer Genome Atlas (TCGA) cohort.

| Characteristics | Summary |
|---|---|
| **Patients with Lung Adenocarcinoma** | N=427 |
| Number of tumor histopathology image series | N=441 |
| Number of histopathology image series of adjacent benign tissue | N=141 |
| Number of histopathology image tiles | N=5,235,656 |
| Age | 65.9 ± 9.9 years |
| Gender | 45.9% Male;<br>54.1% Female |
| Race | |
| White | 291 (68.1 %) |
| Black or African American | 28 (6.55 %) |
| Asian | 5 (1.17 %) |
| American Indian or Alaska Native | 1 (0.23 %) |
| Others | 63 (14.8 %) |
| Unreported | 39 (9.13 %) |
| Anatomic subdivision of neoplasm | |
| L-Upper | 101 (23.7 %) |
| L-Lower | 61 (14.3 %) |
| R-Upper | 136 (31.9 %) |
| R-Middle | 20 (4.68 %) |
| R-Lower | 64 (15.0 %) |
| Bronchial | 1 (0.23 %) |
| Others | 2 (0.47 %) |
| Unavailable | 42 (9.84 %) |
| Stage | |
| Stage IA | 87 (20.4 %) |
| Stage IB | 109 (25.5 %) |

| | |
|---|---|
| Stage IIA | 39 (9.13 %) |
| Stage IIB | 57 (13.3 %) |
| Stage IIIA | 61 (14.3 %) |
| Stage IIIB | 10 (2.34 %) |
| Stage IV | 20 (4.68 %) |
| Stage unavailable | 44 (10.3 %) |
| Grade | |
| Grade 1 | 47 (11.0 %) |
| Grade 1-2 | 9 (2.11 %) |
| Grade 2 | 146 (34.2 %) |
| Grade 2-3 | 28 (6.56 %) |
| Grade 3 | 138 (32.3 %) |
| Grade 4 | 5 (1.17 %) |
| Grade unavailable | 54 (12.6 %) |
| **Patients with Lung Squamous Cell Carcinoma** | N=457 |
| Number of tumor histopathology image series | N=427 |
| Number of histopathology image series of adjacent benign tissue | N=206 |
| Number of histopathology image tiles | N=4,628,373 |
| Age | 67.7 ± 8.5 years |
| Gender | 74.2% Male;<br>25.8% Female |
| Race | |
| White | 301 (65.9 %) |
| Black or African American | 29 (6.35 %) |
| Asian | 9 (1.97 %) |
| American Indian or Alaska Native | 0 (0 %) |
| Others | 114 (24.9 %) |
| Unreported | 4 (0.88 %) |
| Anatomic subdivision of neoplasm | |

| | |
|---|---|
| L-Upper | 122 (26.7 %) |
| L-Lower | 73 (16.0 %) |
| R-Upper | 121 (26.5 %) |
| R-Middle | 17 (3.72 %) |
| R-Lower | 91 (19.9 %) |
| Bronchial | 14 (3.06 %) |
| Others | 9 (1.97 %) |
| Unavailable | 10 (2.19 %) |
| Stage | |
| Stage IA | 79 (17.29 %) |
| Stage IB | 139 (30.42 %) |
| Stage IIA | 60 (13.13 %) |
| Stage IIB | 85 (18.6 %) |
| Stage IIIA | 60 (13.13 %) |
| Stage IIIB | 22 (4.81 %) |
| Stage IV | 7 (1.53 %) |
| Stage unavailable | 5 (1.09 %) |
| Grade | |
| Grade 1 | 9 (1.97 %) |
| Grade 1-2 | 4 (0.88 %) |
| Grade 2 | 168 (36.76 %) |
| Grade 2-3 | 27 (5.91 %) |
| Grade 3 | 197 (43.1 %) |
| Grade 3-4 | 2 (0.44 %) |
| Grade 4 | 9 (1.97 %) |
| Grade unavailable | 41 (8.97 %) |

**Supplemental Table 2.** Characteristics of lung adenocarcinoma and lung squamous cell carcinoma patients in the International Cancer Genome Consortium (ICGC) cohort.

| Characteristics | Summary |
|---|---|
| **Patients with Lung Adenocarcinoma** | N=87 |
| Number of tumor histopathology image series | N=79 |
| Number of histopathology image series of adjacent benign tissue | N=57 |
| Number of histopathology image tiles | N=406,080 |
| Age | 66.5 ± 9.63 years |
| Gender | 47.6% Male;<br>52.4% Female |
| Race | |
| White | 75 (86.2 %) |
| Black or African American | 5 (5.75 %) |
| Asian | 2 (2.30 %) |
| American Indian or Alaska Native | 0 (0 %) |
| Other | 2 (2.30 %) |
| Unreported | 3 (3.45 %) |
| Anatomic subdivision of neoplasm | |
| L-Upper | 16 (18.4 %) |
| L-Lower | 15 (17.2 %) |
| R-Upper | 29 (33.3 %) |
| R-Middle | 4 (4.60 %) |
| R-Lower | 20 (23.0 %) |
| Bronchial | 0 (0.23 %) |
| Unavailable | 3 (3.45 %) |
| Stage | |
| Stage IA | 27 (31.0 %) |
| Stage IB | 24 (27.6 %) |
| Stage IIA | 9 (10.3 %) |

| | |
|---|---|
| Stage IIB | 11 (12.6 %) |
| Stage IIIA | 7 (8.05 %) |
| Stage IIIB | 1 (1.15 %) |
| Stage IV | 5 (5.75 %) |
| Stage unavailable | 3 (3.45 %) |
| Grade | |
| Grade 1 | 12 (13.8 %) |
| Grade 1-2 | 2 (2.30 %) |
| Grade 2 | 29 (33.3 %) |
| Grade 2-3 | 11 (12.6 %) |
| Grade 3 | 29 (33.3 %) |
| Grade 4 | 0 (0 %) |
| Grade unavailable | 4 (4.6 %) |
| **Patients with Lung Squamous Cell Carcinoma** | N=38 |
| Number of tumor histopathology image series | N=33 |
| Number of histopathology image series of adjacent benign tissue | N=19 |
| Number of histopathology image tiles | N=159,272 |
| Age | 71.4 ± 8.0 years |
| Gender | 74.3% Male;<br>25.7% Female |
| Race | |
| White | 33 (86.8 %) |
| Black or African American | 2 (5.26 %) |
| Asian | 0 (0 %) |
| American Indian or Alaska Native | 0 (0 %) |
| Unreported | 3 (7.89 %) |
| Anatomic neoplasm subdivision | |
| L-Upper | 8 (21.1 %) |
| L-Lower | 2 (5.26 %) |

| | |
|---|---|
| R-Upper | 11 (28.9 %) |
| R-Middle | 1 (2.63 %) |
| R-Lower | 13 (34.2 %) |
| Bronchial | 0 (0 %) |
| Other | 0 (0 %) |
| Unavailable | 3 (7.89 %) |
| Stage | |
| Stage IA | 8 (21.1 %) |
| Stage IB | 11 (28.9 %) |
| Stage IIA | 6 (15.8 %) |
| Stage IIB | 5 (13.2 %) |
| Stage IIIA | 3 (7.89 %) |
| Stage IIIB | 0 (0 %) |
| Stage IV | 0 (0 %) |
| Unavailable | 5 (13.2 %) |
| Grade | |
| Grade 1 | 0 (0 %) |
| Grade 1-2 | 0 (0 %) |
| Grade 2 | 18 (47.4 %) |
| Grade 2-3 | 4 (10.5 %) |
| Grade 3 | 15 (39.5 %) |
| Grade 3-4 | 0 (0 %) |
| Grade 4 | 0 (0 %) |
| Grade unavailable | 1 (2.63 %) |

**Supplemental Table 3.** Performance comparison of machine learning methods for lung cancer diagnosis in the TCGA test set. Areas under the receiver operating characteristics curves (AUCs) of the classifiers for distinguishing lung adenocarcinoma (LUAD) from adjacent dense benign tissues, lung squamous cell carcinoma (LUSC) from adjacent dense benign tissues, and LUAD from LUSC are shown. The shaded rows indicate results from our methods, and the unshaded rows indicate results from previously proposed feature-based methods[46].

| **Machine Learning Methods** | **LUAD versus Dense Benign Tissue** | **LUSC versus Dense Benign Tissue** | **LUAD versus LUSC** |
|---|---|---|---|
| AlexNet | 0.94 | 0.98 | 0.90 |
| GoogLeNet | 0.97 | 0.99 | 0.93 |
| VGGNet | 0.97 | 0.98 | 0.93 |
| ResNet | 0.95 | 0.94 | 0.88 |
| SVM with Gaussian Kernel | 0.85 | 0.88 | 0.75 |
| SVM with Linear Kernel | 0.82 | 0.86 | 0.70 |
| SVM with Polynomial Kernel | 0.77 | 0.84 | 0.74 |
| Naïve Bayes Classifiers | 0.73 | 0.77 | 0.63 |
| Bagging | 0.83 | 0.87 | 0.74 |
| Random Forest using Conditional Inference Trees | 0.85 | 0.87 | 0.73 |
| Breiman's Random Forest | 0.85 | 0.87 | 0.75 |

**Supplemental Table 4.** Transcriptomic subtype of lung adenocarcinoma and lung squamous cell carcinoma patients with available histopathology slide and RNA-sequencing data.

| **Lung Adenocarcinoma** | |
|---|---|
| Terminal Respiratory Unit (TRU) | 172 (35.98 %) |
| Proximal Inflammatory (PI) | 168 (35.14 %) |
| Proximal Proliferative (PP) | 138 (28.87 %) |
| **Lung Squamous Cell Carcinoma** | |
| Classical | 165 (33.88 %) |
| Basal | 133 (27.31 %) |
| Secretory | 119 (24.43 %) |
| Primitive | 70 (14.37 %) |
